# Trends in Coupled Human-Environment Systems Modelling: A Scoping Review

**DOI:** 10.1101/2024.11.08.618889

**Authors:** Vivek A. Thampi, Chris T. Bauch, Madhur Anand

## Abstract

Classical environmental models assume the influence of humans on environmental systems is constant. However, human and environmental systems respond to one another. As such, coupled human-environment systems (CHES) models have been developed and are becoming more widely studied. In this review, we analyze CHES modelling techniques and study systems over a decade (May 2009-April 2019). We utilized the PRISMA method to filter publications from both Web of Knowledge and PUBMED, yielding 92 relevant papers for our review. Publications more than doubled from the 5-year interval May 2009-December 2013 (28/92) to the 5-year interval January 2014-April 2019 (64/92). CHES models typically used either differential equations (DEs) (44/92) or agent-based models (ABMs) (28/92). We organized the included literature with respect to the technique used to represent human behaviour. We noticed a diversity of approaches in this respect, but primarily optimization techniques (28/92) and game theory (34/92). We noticed a substantial increase in publications using more highly structured models in the second 5-year interval. We attribute this to reduced technological barriers to developing more detailed models, and greater data availability. We discuss the realism of the models and their ability to capture real-world dynamics. Finally, we explore avenues for future research, and discuss unconventional routes such as online communities and artificial intelligence modelling to expand representation of human behaviour in CHES models.

## 1 Introduction

Ecosystem sustainability and human impact on ecosystem dynamics have been a focus of environmental research due to the importance of ecosystem services in various sectors such as agriculture, fisheries, and tourism, as well as support for natural ecosystem conservation. Ecosystems have undergone deterioration through various sources of disturbance. Damage caused by natural events, such as forest fires in fire-prone regions, can cause significant shifts in ecosystem dynamics if the burns become unnaturally large or frequent. In some cases ecosystems are sufficiently resilient to bounce back from low-impact natural disruptions. This is exemplified by the *Diadema antillarum* sea urchins consuming algal turfs to promote new coral establishment [1], or seed dispersal during the aforementioned fires to mitigate the damage and loss caused by the forest fires [97].

Humans can also have negative impacts on ecosystem services, causing a decline in resilience and often catastrophic disruptions. These disruptions can take many forms, such as eutrophication of lakes via pollution [42] or depletion of a natural resource via exploitation. These disruptions may not always be intentional. For instance, travelling individuals have unintentionally introduced pests or exotic species to forest ecosystems. This has occurred through individuals importing pests via infested firewood [12], or through exotic plant growth, which introduces competitive interactions into grassland dynamics [112]. The resulting regime shifts have often been characterized by a hysteretic feedback loop, wherein greater effort is required to reverse the catastrophic trajectory an ecosystem has taken, than what caused the catastrophic shift in the first place.

Human interventions are not always negative, however. Research has explored how conservationist strategies can emerge in response to disturbances to promote sustainability or recovery in affected systems [50, 102]. These efforts can range from the development of sanctuaries for protecting endangered species, or raising awareness of personal ecological footprints in a population[113]. These strategies, however, may not propagate through a population, if individuals are dissuaded from these strategies by incentives to act unsustainably. Therefore interventions seek ways to incentivize individuals to opt into acting in the interest of the community as a whole [40]. By promoting the development of social norms (peer influence) and social learning (fostering community and online interactions), individuals can be influenced to modify their strategies by changing their perceived utility in a way that benefits ecosystem integrity.

Environment system modelling has played a strong role in both developing and shaping the strategies required to promote ecological resilience and recovery. Classically, modelling techniques have incorporated fixed parameters to represent human input in ecological models (and vice versa)[96, 53]. However, human systems and ecosystems are rarely constant, and feedback between the systems, such as those mentioned above, can influence human and/or environmental behaviour. Such feedbacks have been observed in various scenarios, such as COVID-19’s effects on social gatherings, or environmental activism sufficiently pressuring industries to adopt more ecologically sustainable alternative strategies. Thus, there is a need to develop models that capture the dynamic nature of human and environment interactions.

Coupled human-environment models incorporate a dynamic coupling between human and environment systems. These coupled human-environment systems (CHES) have been developed to incorporate a two-way feedback mechanism that integrates responses from both human and environment dynamics in a cyclical manner (Figure 1). As observed, this feedback loop represents the non-static nature of human behaviour, where strategies can be changed to optimize an individual’s utility (i.e., subjective preferences). This methodology is often more realistic compared to static human parameter representations in classical environmental system models and has been utilized in a variety of ways in environmental systems research.

**Figure 1:**
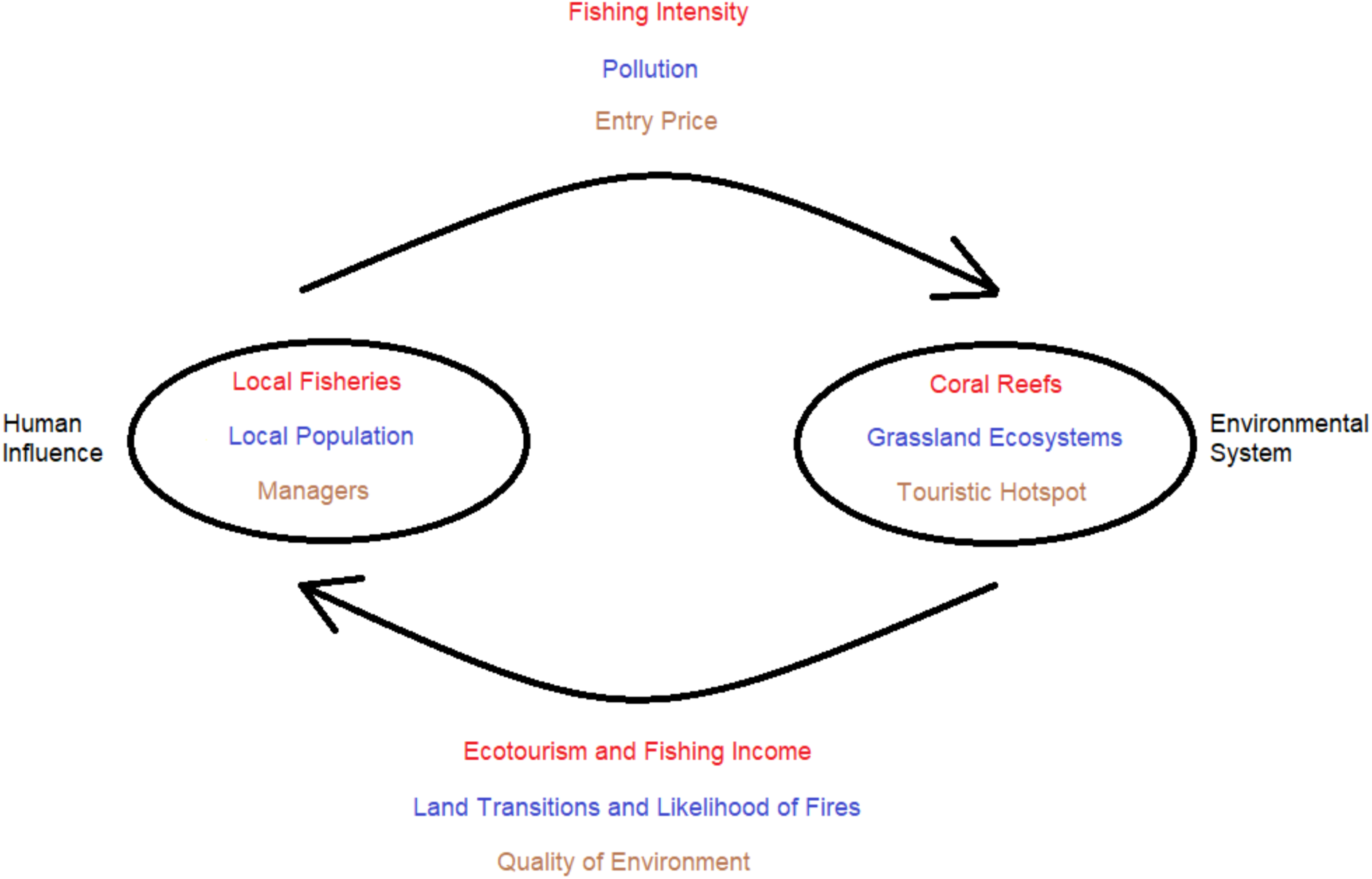
Cyclic nature of human behaviour to current state of an observed environmental system that is modelled using coupled human-environment systems.

Numerous modelling methodologies have been developed that incorporate a dynamic link between human and environmental components and have ranged from extensions of classical modelling techniques (such as the Lotka-Volterra competition/fishery models) [96, 53] to more complex Cellular Automata-Markov chain models that investigate stochastically-driven land state transitions [118]. In addition, very complicated and detailed computational models and tools have also been developed [10, 7]. Typically, these modelling techniques require numerous input parameters for both environmental and human components, and as such can be computationally expensive. With technological improvements, computational models have become more feasible, enhancing the viability of these modelling techniques.

Human behaviour can be incorporated using a game-theoretic approach. This methodology assumes that individuals act to maximize their payoff (or utility). Individuals choose strategies from a strategy set, and their payoff for a given strategy depends on what strategies other players adopt [110]. As an example, we consider the two-player Prisoner’s Dilemma [5]. The game supposes two individuals who have been caught under suspicion of committing a crime and are being interrogated in separate cells. A player may choose to cooperate with the other player (by not confessing), or defect against the other player (by confessing to the crime in the hope of a shortened prison sentence). If both players cooperate, they avoid a prison sentence. If both players defect, they suffer a shortened sentence. If one player cooperates and the other defects, the defector gets an even shorter prison sentence, while the cooperator gets a very long prison sentence. The prisoners would obtain the best outcome if they both cooperated, but game theory predicts that defection will be the strategy they both choose. This simple game illustrates the clash between what is socially optimal versus the individually optimal actions that individuals adopt in practice. This clash between socially and individually optimal outcomes occurs in many common pool resource problems in the environmental sciences. In more sophisticated games, payoff functions can be tailored to represent various strategies in a given ecological scenario. CHES in this context can be thought of as systems where individuals behave according to utility functions that depend on environmental states, which respond to the strategy choices of other members of the population, in turn.

To our knowledge, no one has performed a recent scoping review of CHES models of ecological and environmental systems. Here, we report a scoping review of the literature on CHES models over a ten year period from May 2009 to April 2019. Our aims are to assess the modelling methodologies, study systems, number of models published over time, and means of implementation of human-environment coupling. We performed the Preferred Reporting Items for Systematic Reviews and Meta-Analyses (PRISMA) methodology to synthesize and filter the literature. We end with a discussion of the future potential of these modelling techniques for capturing complex dynamics of human and environment interactions.

## 2 Methodology

### 2.1 Methods Overview

In the following sections, we outline the process used to perform our scoping review. Searchterms were developed to capture papers on coupled human-environment system models. Searches used both Web of Knowledge (WOK) and PUBMED databases to generate a collection literature organized by titles, abstracts and keywords over a ten-year period. These results were synthesized using the PRISMA process (detailed below) to identify modelling research that utilized coupled human-environment systems.

### 2.2 Necessary Criteria

Qualifying research literature were required to satisfy the following criteria:

1. *An Environmental or Ecological Foundation*: Collected literature was required to have a focus on environmental or ecological systems. These included topics of sustainability, ecological management, water allocation, tourism, etc. We included models that were not developed TOPIC: (((”social and ecological” OR “ecological and social” OR “human-environment” OR “socioecological” OR “human-and-natural” OR “CHANS” OR “human and environment” OR “human-natural” OR “socio-ecological” OR “Natural-Human” OR “environmental-Human” OR “Environment-Human” OR “social-ecological”) AND (”mathematical model*” OR “differential equation*” OR “simulation model*” OR “dynamic model*” OR “compartment model*” OR “system dynamics” OR “Markov Chain*” OR “Generalized Modeling” OR “Generalized Modelling” OR “Decision Model” OR “Theoretical Model” OR “Decision-Making Model”))) OR TOPIC: ((”human-environment system model” OR “human-environment model” OR “Human-environment dynamics” OR “socio-ecological system model” OR “socioecological model” OR “socio-ecological dynamics” OR “human-and-natural system model” OR “human-and-natural system dynamics” OR “CHANS model” OR “ecological and social dynamics” OR “social and ecological dynamics” OR “environmental-human system*”)) for a specific system but rather represented some general class of ecological or environmental systems (we refer to these as ‘general system models’).
2. *Mathematical Modelling Techniques*: We aimed to collect literature that utilized mathematical modelling, such as systems of differential equations. These can also include computational simulations. Models were required to exhibit coupling between human and environment systems. Examples ranged from land-use land change (LULC) models, to cellular automaton (CA)-Markov models. This condition also includes simulating population with artificial intelligence or machine learning techniques, and can include game-theoretic ecological modelling techniques.

### 2.3 Exclusions

To exclude irrelevant papers and limit the scope of the review, we developed a preliminary screening criteria for the PRISMA process which targeted and removed results from epidemiology (vaccinations, disease dynamics, etc.), urbanization, industrial ecology (work safety, worker-ecosystem, etc.), and other non-environmental related work.

### 2.4 Search Terms

The search terms are provided in Table 1.

**Table 1:**
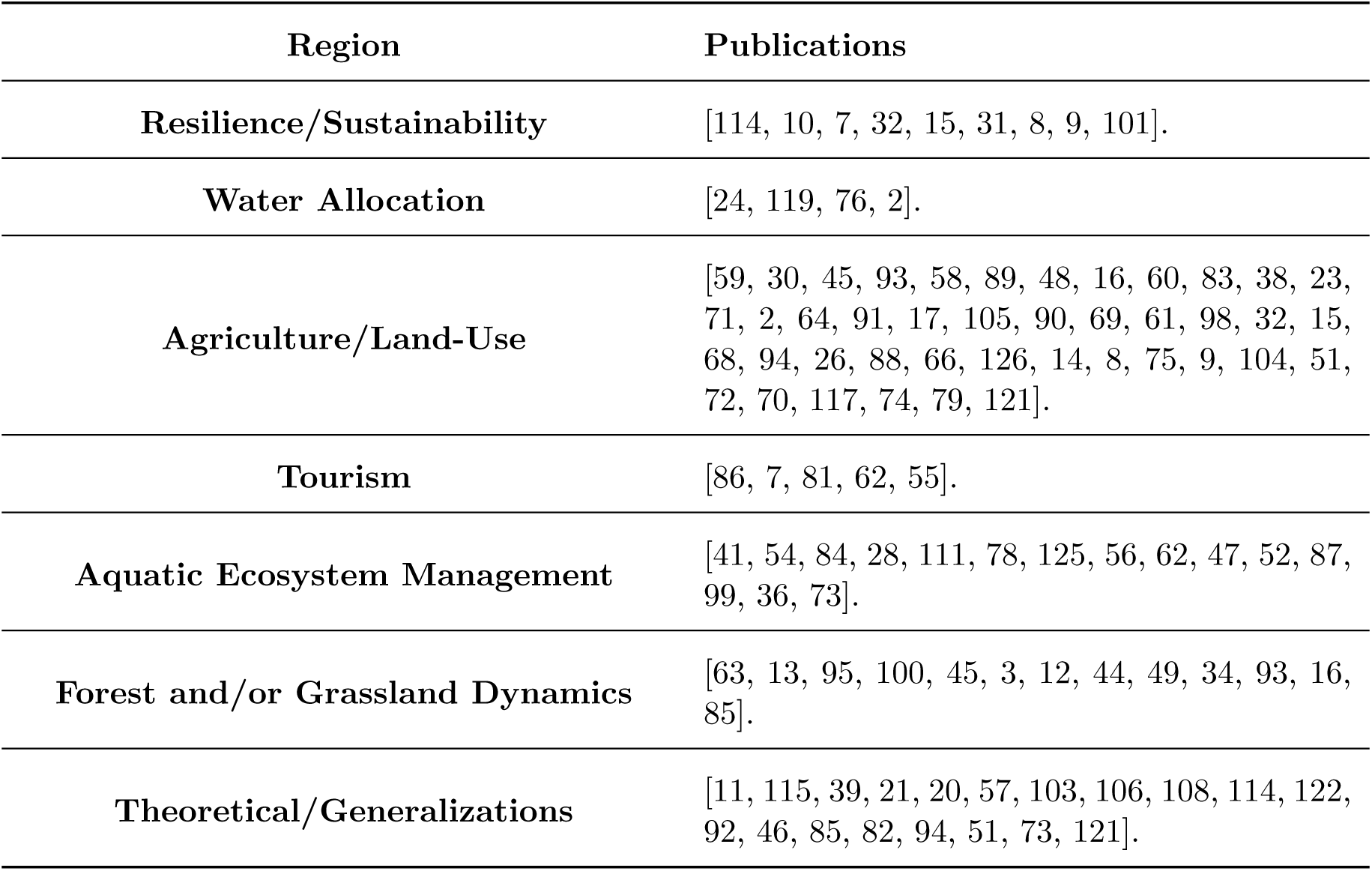
Search terms that were inputted into both Web of Knowledge, and PUBMED databases between May 2009 to April 2019.

### 2.5 PRISMA

To identify relevant literature for this review, we used PRISMA, a tool that has been established since 2018 useful for systematic scoping reviews. PRISMA uses a step-by-step methodology to identify the most relevant papers from any collection of research [80]. The process and the number of papers we identified at each step of the PRISMA method are:

Step 1: Papers were collected using the search terms in Table 1 using Web of Knowledge and PubMed. Search results were saved using containing title, author, abstract and keywords. 2188 papers were obtained in total.

Step 2: Papers were assessed manually, filtering out any duplicates that were obtained from both search engines. This decreased the number of relevant papers to 1922.

Step 3: Papers were further screened based on their relevance to the review topic after scanning the titles, keywords and abstracts. Models needed to pertain to environmental/ecological systems and needed to have some mention a related human component which impacts the modelled system dynamics. The number of relevant papers decreased further to 681.

Step 4: Papers were further screened after a more thorough reading of the obtained literature. During this process, original research were retained while reviews and papers unrelated to Ecology or Environmental Science were removed. A total of 92 papers remained upon the completion of PRISMA.

## 3 Results

### 3.1 Notable Highlights

Several temporal trends and patterns in the collected literature were clear. Nearly 33% of reviewed publications applied some form of real-world data (questionnaries, surveys, etc.) for use in model parameterization or calibration. Furthermore, a large subset of publications developed models for a generalized ecological system (19/92) rather than for a specific ecosystem type. Lastly, the largest subset of publications focused on agricultural settings (21/92), while the smallest subset focused on climate change dynamics (2/92).

### 3.2 Date of Publication

Results were organized based on publication date. The number of publications over the time window of the review showed variability from year to year, but overall they show an increasing trend (Figure 2). It should be noted that the endpoints do not accurately describe the total number of papers in those specific years since the review time window started in the month of May and ended in the month of April.

**Figure 2:**
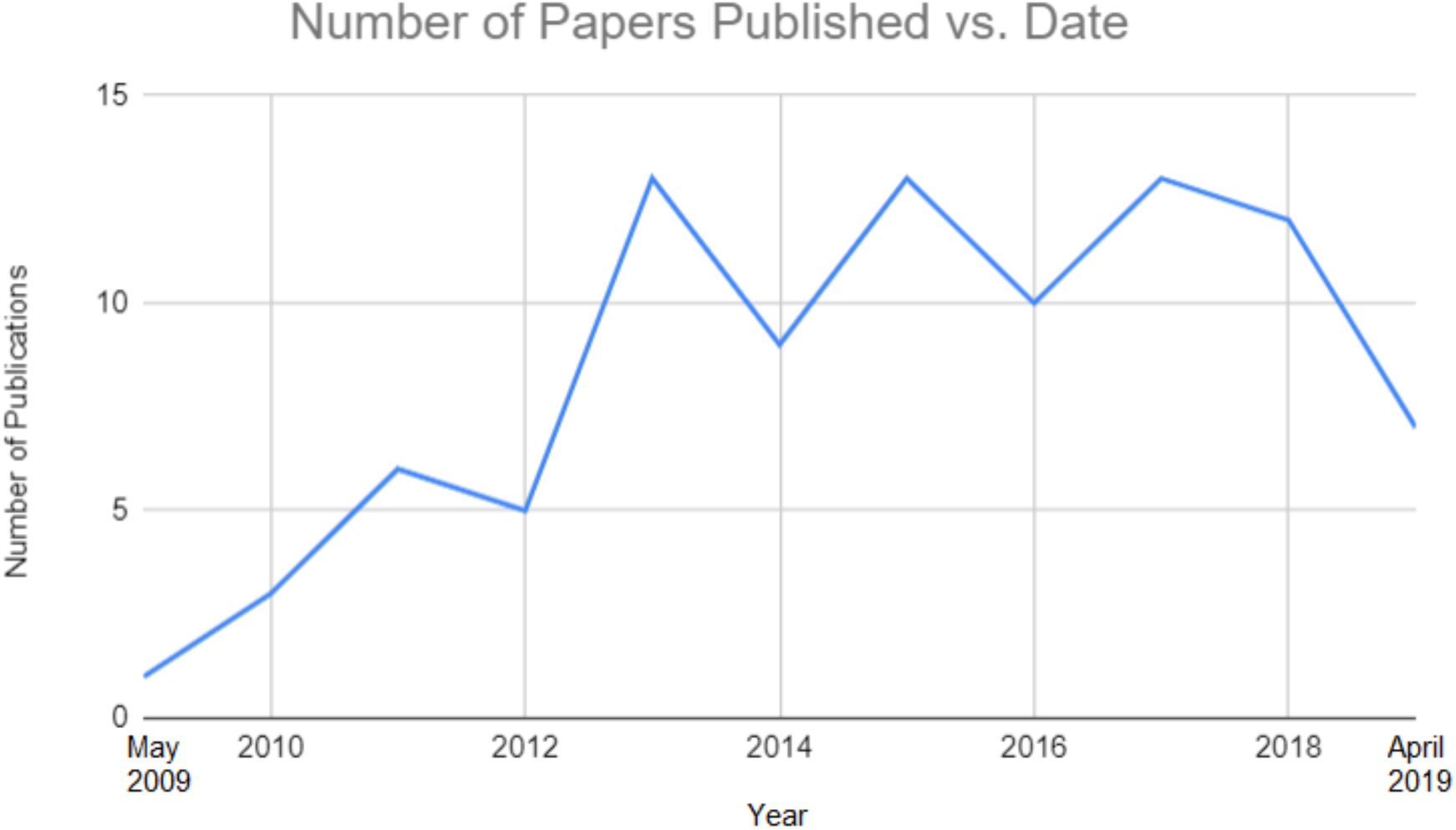
Literature organized by publication year. Papers were published between May 2009 and April 2019. Hence, the number of publications plotted for 2009 and 2019 only represent a subset of papers published in those calendar years.

### 3.3 Environmental System Topics

Results were also organized by the modelled system. Most papers focused on agricultural systems (41/92), and especially optimization of pastoral land-use benefits and land use transitions. We separated water allocation (4/92) from both land-use models and aquatic ecosystem management, since the topics of papers studying aquatic ecosystems (15/92) ranged from ecosystem well-being to optimization of fisherman benefits. Topics concerning tourism (5/92) were classified separately, since touristic CHES models were sufficiently numerous, regardless of the ecological setting they modelled. Forests and grassland models were combined under one classification (13/92), as were topics focusing solely on resilience or sustainability (9/92). Finally, many CHES models were general system models, developed without reference to a particular study system (19/92). These models were included due to their applicability to different environmental topics that were usually mentioned in the paper’s Discussion section. Results have been compiled in Table 2.

**Table 2:**
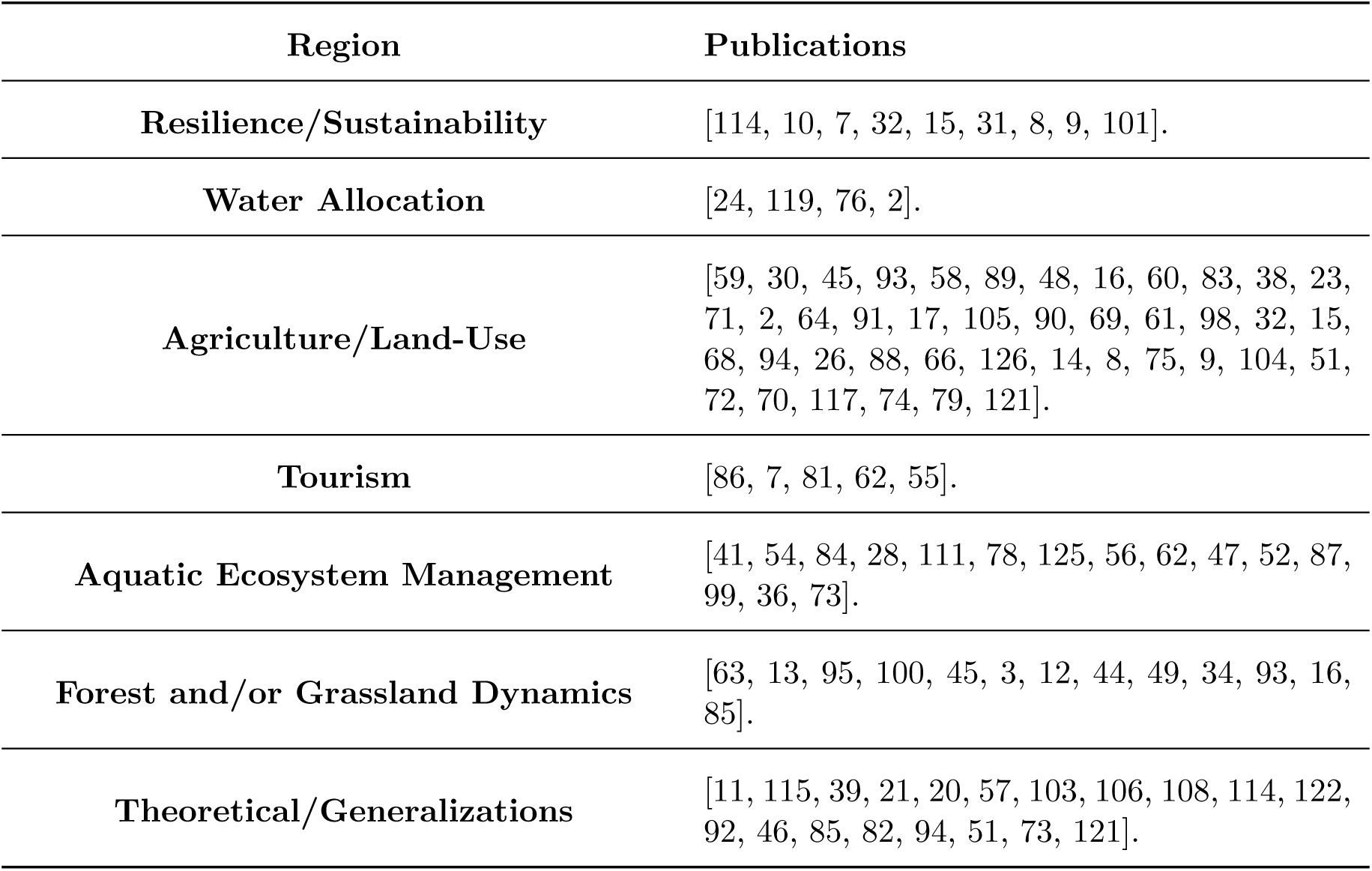
Publications organized by study.

### 3.4 Model Type

Results were categorized by their model type to highlight the various methods of representing coupled human-environment systems. Models were placed into four different classifications Differential equations (DEs), Agent (Individual)-Based Modelling (ABM/IBM), non-ABM stochastic models, and unspecified/other models, which are discussed in more detail below. Publications by model type are summarized in Table 3. We also plotted the number of each model type in each year. We noticed steady growth of DE models until 2016 following a decline, while the number of ABMs varied throughout the review time period. The number of stochastic models each year was roughly constant, while unspecified/other modelling techniques grew in the latter half of the review time period. The results were compiled and can be seen in Figure 3.

**Figure 3:**
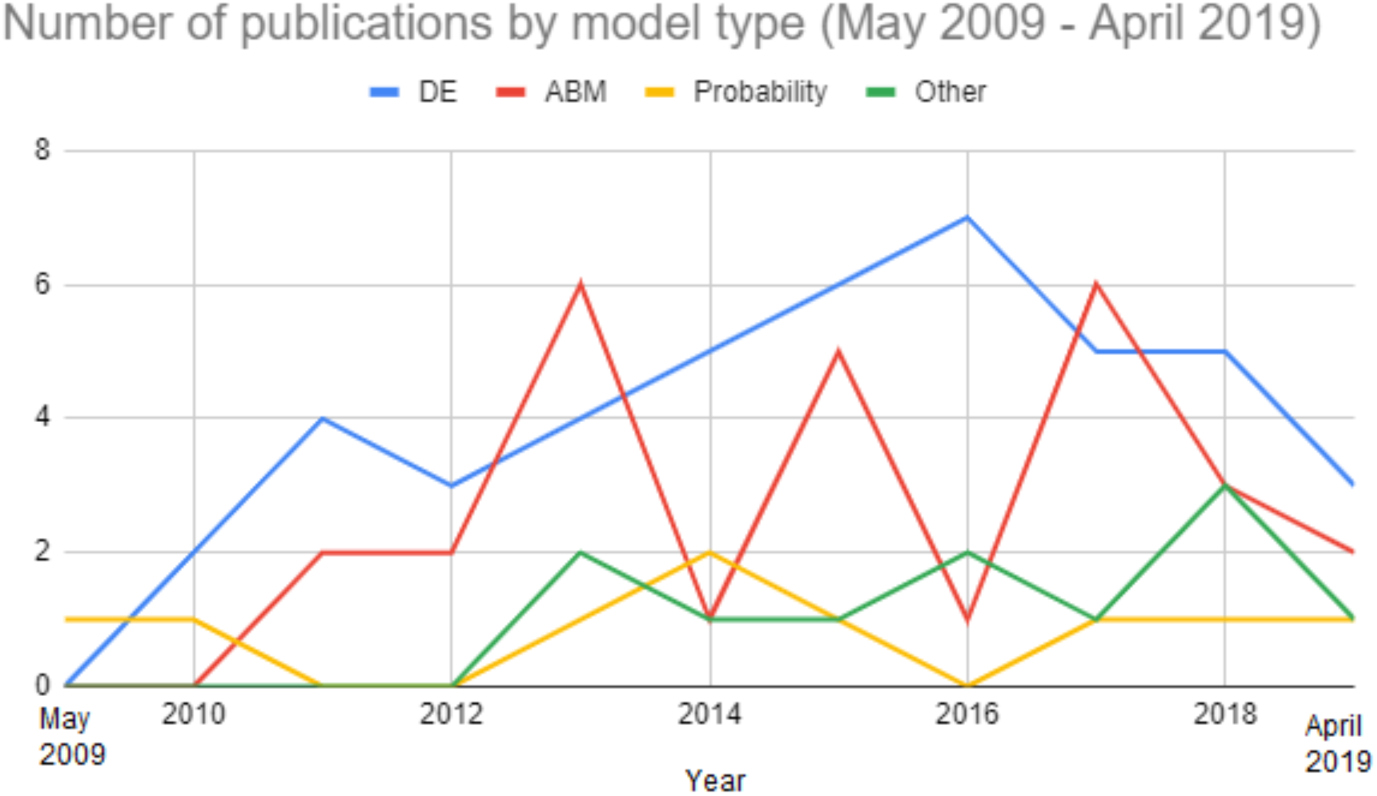
Number of publications using different model types between May 2009-April 2019. Because the start and end dates of our study fell partway through May 2009 and April 2019, the number of publications plotted for those years do not represent the entire calendar year.

**Table 3:** Publications organized by model type.

| **Model Type** | **Publications** |
| --- | --- |
| **Difference/Differential Equations** | [59, 63, 115, 84, 13, 39, 28, 21, 20, 95, 111, 100, 45, 78, 125, 3, 56, 12, 49, 57, 34, 58, 103, 114, 48, 38, 81, 71, 64, 17, 69, 62, 47, 46, 55, 52, 85, 31, 94, 75, 51, 99, 101, 73]. |
| **Agent(/Individual) - Based Modelling** | [11, 54, 30, 86, 89, 108, 76, 60, 16, 83, 23, 6, 2, 91, 92, 105, 90, 61, 98, 82, 68, 126, 14, 104, 70, 117, 36, 79]. |
| **Stochastic Models** | [44, 93, 106, 119, 122, 32, 88, 66, 121]. |
| **Other** | [41, 24, 10, 7, 87, 15, 26, 8, 9, 72, 74]. |

#### 3.4.1 Differential Equations

Differential equations (DE) have provided the foundation for much mathematical modelling. Differential equations describe how system variables evolve dynamically in response to one another as a function of model parameters in continuous time. Differential equations can capture behaviour near equilibrium states, transient dynamics, and sudden transformations–regime shifts–in a natural way. In the case of classical mathematical modelling of environmental systems, these models represented human influence implicitly by considering the impact of humans on natural system parameters such as fecundity or flow rates, whereas human-environment models represent the human system explicitly–as a state variable–that itself is influenced by other parameters such as mitigation cost, for instance.

Differential equation models of natural systems have been expanded into CHES models to explore a number of topics in human-affected ecosystems such as natural land depletion taxation [59], incentivization of fisherman behaviour [84], or conservationist opinion and behaviour propagation in populations through social processes [13].

Deterministic differential equations are a subset of DEs where the model trajectory is determined entirely by parameter values and initial conditions. Deterministic CHES DE models vary in complexity and implementation. In [49], Innes et al, conservation opinions regarding Brazilian forest-grassland mosaic model evolved dynamically according to ecosystem rarity. The perceived utility of individuals influenced land-use decision making. Horan et al [46] utilized decision dynamics in a modified Lotka-Volterra predator-prey model to investigate harvesting dynamics on crayfish and bass biomasses. In order to develop the human component of the model, fisherman harvesting choices were conditioned on the relative costs, benefits and regulation policies. In both models, behavioural components dynamically affected environmental system dynamics and *vice versa*.

An alternative DE methodology was developed to investigate the effects of mass tourism versus ecotourism by Monfared et al. [81]. The DE model was composed of a 4-dimensional dynamical system to investigate the ecological impact between mass tourists and ecotourists. In their model, two compartments were developed to represent different members of the population mass tourists and eco-tourists, and another two compartments to asses changes in capital and ecosystem quality. Mass tourists exploit the ecosystem services regardless of its current status, whereas eco-tourists refrain from visiting these sites when environment system dynamics cannot adequately sustain visitors. Interactions between both tourists (similar to mass-action mixing components in disease models) would dissuade both populations of tourists from visiting. Coexistence of both different populations, albeit chaotic in nature, was proven possible through numerical analysis, and would lead to unsustainable (oscillatory) environmental quality.

Stochastic differential equations constitute another subset of DE models. These models incorporate stochasticity (’random noise’) processes. An example of these models was developed by Ali et al [3] where a stochastic coupled human-environment system of pest invasions in provincial parks was developed. Imitation dynamics were observed to drive changes between strategies of buying firewood locally or transporting it. Social norms were incorporated to influence strategy switching under the concern of potential infestation. Another stochastic system developed by Lee et al [64], coupled a dynamic model of grassland dynamics to foraging decision-making. They used stochastic best-response dynamics, which is a modified technique of classic best-response dynamics where players choose the optimal outcome at the next time step, subject to a stochastic term that represents other influences on their decision-making process [35]. Bistability, where a system may end up either in one state or a different state, depending on initial conditions, was also observed. Furthermore, model dynamics admitted oscillatory behaviour between relative grass biomass and herder populations, where depletion of grasses causes herders to vacate the area. Given sufficient system resource resiliency, depleted grasses are able to recover over time and maintain the cyclic relationship between grass biomass availability.

Some papers used custom-built software to develop and analyze differential equation models. In Marin et al [69], a CHES DE model was developed with a mathematical modelling and simulation software package called STELLA, to investigate the impacts of social capital on scrubland dynamics. Another CHES model was developed by McClanahan et al. [78] in STELLA to investigate the dynamics of restrictions and regulations on coral reef fisheries. This software package uses compartmental diagrams to provide insight and aid in determining the appropriate linking elements in the model. Once all required elements of the model are defined, simulations can be generated and tested with empirical data [65].

#### 3.4.2 Agent-Based/Individual-Based Modelling

Coupled human-environment systems can also be represented using agent-based modelling (ABM). These models employ autonomous agents to perform pre-defined actions that represent the role that they have been assigned [116]. Agents are capable of interacting with other agents, based on the design criteria of these models. These agents can also perform various decision-making processes based on the status of the system, sometimes in order achieve an optimal solution set, while taking into account factors such as demand and other predefined criteria. These can be visualized as pathway-based systems which performs actions upon performing an assessment of all collected possibilities and their overall payoffs [19].

An example of these models was used by Berfuss et al [11] who developed an ABM coupled human-environment system to explore policy making and its effects on economic growth with applications to climate change and fisheries. The model used Markov decision making processes, which processes decisions partially by choice and partially randomly, to characterize agent behaviour. Triggers were found to influence the adoption of both unsafe and unsustainable policy making strategies if agents blindly followed an economic optimization strategy. Another coupled human-environment system developed by Coutts et al [30] utilized an ABM to investigate agent behaviour in response to weed spread in pastoral regions. Agents relied on factors such as profit, invasion probability and social pressures when performing decision making processes. Weed prevalence was directly influenced via social pressure. This was also in part due to the effect of perceived benefits on how internal social norms influenced the severity of the weed prevalence.

Similar to DEs, software packages that allow users to develop and analyze ABMs have been used. Work by Synes et al [108] developed an ABM CHES to create a competitive scenario where social agents compete for land based on their ability to effectively use capital and ecosystem services. Modelling processes were simulated with the computational program CRAFTY, using ecosystem service levels to provide the basis for agent decision-making processes with respect to land-management. Results indicated a need to use agent-functional types of models to facilitate ABM application across a large spatial extent, and a potential loss of information by employing uncoupled modelling techniques. In addition, another CHES ABM model was developed Marohn et al [70] that coupled two software packages LUCIA (soil, water and plant dynamics) and MP-MAS (farmland decision making, investment, production and income optimization). Their model assessed conservation strategies in a highland agricultural setting. They identified low-cost conservation strategies that could benefit soil quality, household income, and farmland productivity. In particular, changes in fertilizer prices can strongly influence agent decisions in choosing to adopt soil conservation practices, and can instead support the choice to maximize crop yield.

#### 3.4.3 Stochastic Modelling

CHES models have also used stochastic modelling techniques such as Markov Chain modelling. Models in this category capture state transitions via a matrix where each element of the matrix represents the probability of a transition from one system state to another one. These can be used in tandem with other modelling techniques or also commonly with CA cellular automata, which evolve based on both a set of predefined rules and the states or results of neighbouring strategies, or using system dynamics, which captures feedback loops and transitions based on predefined probabilities. Transition matrices are developed to illustrate the probability of an agent’s decision and their impact on natural system states.

Henderson et al [44] illustrate this methodology with a Markov chain model of landowner decisionmaking based on preferences and land cover in each time step. Individuals may choose to clear a given patch of forest or have it harvested for lumber in each time step. Using utility-based decision making, it was determined that implementing certain conservation incentives did not support stable forest cover, and instead caused cyclical bouts of deforestation due to interactions between individual landowner decisions and total amount of forest cover and harvesting decisions in the whole population.

### 3.5 Human Component

The methods used to represent the human component of CHES varied widely in the included literature (Table 4). We explore these various approaches in the following paragraphs.

**Table 4:**
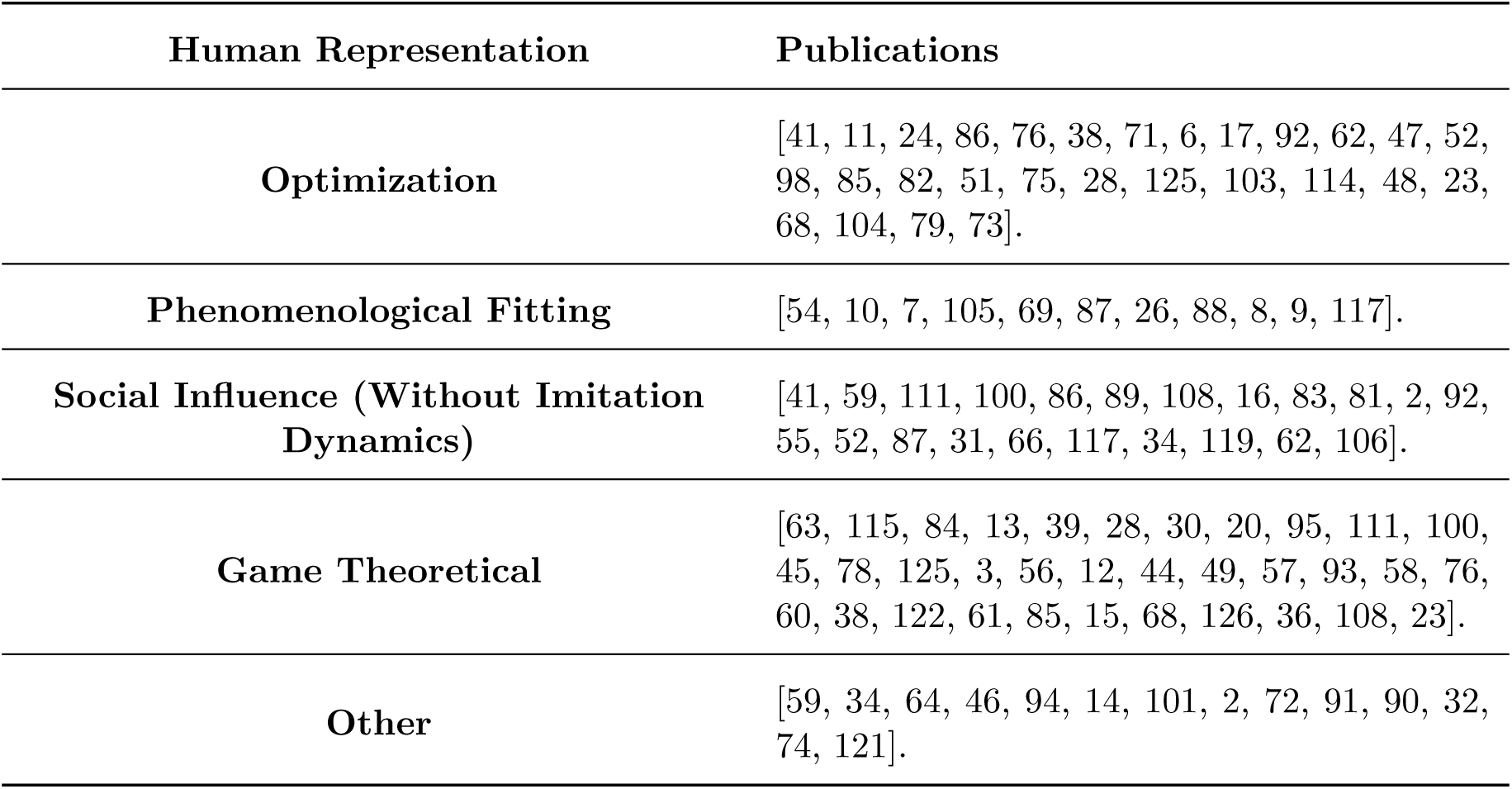
Publications organized by the methods used to represent the human system.

| Human Representation | Publications |
| --- | --- |
| **Optimization** | [41, 11, 24, 86, 76, 38, 71, 6, 17, 92, 62, 47, 52, 98, 85, 82, 51, 75, 28, 125, 103, 114, 48, 23, 68, 104, 79, 73]. |
| **Phenomenological Fitting** | [54, 10, 7, 105, 69, 87, 26, 88, 8, 9, 117]. |
| **Social Influence (Without Imitation Dynamics)** | [41, 59, 111, 100, 86, 89, 108, 16, 83, 81, 2, 92, 55, 52, 87, 31, 66, 117, 34, 119, 62, 106]. |
| **Game Theoretical** | [63, 115, 84, 13, 39, 28, 30, 20, 95, 111, 100, 45, 78, 125, 3, 56, 12, 44, 49, 57, 93, 58, 76, 60, 38, 122, 61, 85, 15, 68, 126, 36, 108, 23]. |
| **Other** | [59, 34, 64, 46, 94, 14, 101, 2, 72, 91, 90, 32, 74, 121]. |

#### 3.5.1 Optimization

Human behaviour was most commonly represented in coupled human-environment systems via the use of optimization techniques. In general, papers in this category optimized functional representations of individuals’ personal interest in a natural system, and predicted the impact of optimized personal decision-making on natural system states. (This approach we treated as distinct from game theory, which will be discussed separately later, where individuals are assumed to optimize their payoff in a situation where their payoff is dependent on the actions of other individuals of the population.) Using a functional form to represent payoffs or utilities, members of the population can be modelled implementing strategy switching (imitation) to follow strategies with the greatest payoff value.

An example of representing human decision-making as optimization of an objective function is exhibited by Blanco-Gutierrez et al [17]. The authors develop an integrated economic-hydrologic CHES model where farmers aimed to maximize their income by assessing three different model scenarios: business-as-usual (BAU), EU-policy driven, and national policy driven scenarios. Results revealed farmers’ behaviour when handling risk, causing the model to predict a tradeoff between risk-taking and overall profit obtained by farmers. Model dynamics revealed detrimental effects of BAU-scenarios, but showed management changes when both EU and national policy measures were implemented.

In a fishery model by Hunt et al [47], a CHES model was developed using a “utility-theoretic” approach to model fisherman angler behaviour. In order to maximize utility, anglers considered a given set of lakes and chose a lake with the highest perceived utility. Model analysis revealed that low angler population sizes and high catch importance reduced the potential for anglers to overfish.

#### 3.5.2 Phenomenological Modelling

Phenomenological models describe the observed dynamics of a system without articulating a mechanistic basis for the components of the model structure. In general, these models utilize empirical data from various sources to establish functional representations for their components. Additionally, in some cases where empirical data is insufficient, parameters can be calibrated based on historically observed trends or biologically plausible trajectories [3]. Examples of phenomenological fitting in the reviewed literature used surveys or other empirical data such as fisherman catch per unit effort (CPUE) or firewood transport costs in order to quantify model parameters governing phenomenologically-justified functions [52, 12].

#### 3.5.3 Artificial Intelligence (AI) and Machine Learning (ML)

AI and ML models have also been developed to simulate the dynamics of human interactions with environmental systems [121]. These approaches are often used to represent human behaviour in a CHES model. But in other cases, investigators have hybridized the use of machine learning algorithms to inform land use modelling, with role-playing games involving real human participants. For instance, Washington-Ottombre et al. [121] use this approach to study land use decisions in a fictional land named Mageria, based on interactions between pastoralists, farmers, and a land commissioner. AI and ML methods were relatively unexplored by the CHES modelling literature, but do possess potential application to capture individual behaviour.

#### 3.5.4 Game Theoretical Techniques

As mentioned previously, game theory can be used to model how humans behave in strategic decision-making where their payoff depends on the strategies chosen by both themselves and other individuals. These methods have also been used in CHES models. One such method employs replicator dynamics, in which individuals do not automatically switch to the highest-payoff strategy at some point in time but rather only switch after some kind of learning or imitation process. For instance, in imitation dynamics, an individual samples other individuals in the population and, if the focal individual encounters someone playing a different strategy, the focal individual switches their strategy with a probability proportional to the perceived payoff gain for switching to that strategy. These methods are implemented using differential equations to develop representative utility/payoff functions that become incorporated into the overall model structure for dynamic human behaviour.

An example of replicator dynamics was observed in a CHES model developed by Henderson et al [45], investigating the effects of coupling human dynamics of land preference on the Brazilian forest-grassland mosaic. Imitation dynamics were developed to capture the preferences of the population when choosing whether to modify or keep land for either agricultural, forest or grassland uses. With modifications to the utilities for the protectionist strategy, the model predicted significant impacts on human-environment stability. Some examples of these effects include high conservation values tending to transform land away from agricultural purposes. Model predictions demonstrated a straightforward response to increasing conservation values of grasslands. Increasing forest conservation values instead caused model dynamics to stabilize to a state governed by forest-grassland limit cycles, or grassland stability. Additionally, with sufficiently high economic discount rates, compared to conservation discount rates, model dynamics tend to stabilize in either forest or grassland states (and conversely stabilize in an agricultural state for sufficiently low economic discount rates).

Lade et al [57] utilized a game-theoretic CHES model to analyze common-pool resource use dynamics. In their work, the human component followed replicator dynamics and payoff functions for defectors and cooperators were defined. Results revealed significant resource depletion driven by humans. In addition, with a constant inflow of a resource, the net benefit of the defection strategy would outweigh the value of a co-operative regulatory strategy, leading to a catastrophic collapse of resource. Given that regime shifts can occur unexpectedly in response to slowly changing external drivers, this work emphasizes the danger of exploiting environmental resources, especially under lack of proper monitoring. The authors emphasize that including human dynamics in their model revealed predictions that would not occur without the dynamic coupling between human and natural systems.

In a model by Rodrigues et al. [93], the impact of human behaviour was implemented in a 2-strategy, 2-person game CHES model of forestry dynamics. In their work, deforestation was utility-driven, with state transitions being described using Markov-chain transition probabilities. Results showed that deforestation would benefit the landowners, but would cause degradation of ecosystem services and value of neighbouring land, especially if forest recovery is sufficiently slow. Overall, forest regeneration would favour mutual cooperative strategies, but would also govern the rate of agricultural abandonment as the decision to conserve would be dependent on the choices of neighbouring landowners. Alternatively, if forest recovery is sufficiently fast, landowners would be more likely to choose to deforest, a similar scenario observed by Lade et al. [34] with resource exploitation. This prediction is an excellent example of the ‘law of unintended consequences’ and demonstrates the value of including feedback between human and natural systems via CHES models.

In a model developed by Mason et al [76], an agent-based evolutionary game model was developed to determine operator decision criteria with respect to water resource systems. In their model, operators (agents) undergo a negotiation process until a successful proposal is achieved. Agents aim to increase their own utility for personal benefit, which would be equivalent to a decrease in utility for unsuccessful agents. Knowledge of the current state would influence future decisions as well as proposals agents make under various climactic conditions. While new proposals cannot directly undermine other agents, new solutions would add to the agent’s total set of proposals, and agents must grant a concession to another agent. Model dynamics were tested on synthetic flood control and water supply dynamics and were successfully able to capture operator behaviour under climatic conditions of extreme wet and dry scenarios.

#### 3.5.5 Social Learning (Without Replicator Dynamics)

A number of CHES models represent social learning and other social processes without invoking replicator dynamics. In these models, in any given population the effects of social influence or interactions can influence individuals to modify their personal strategies in order to maximize their personal gain, without using the specific structure of the replicator equations. Rebaudo and Dangles [89] developed a CHES model with a human component that utilized information transferal mechanisms to investigate the role of social dynamics in pest dynamics. In their model, agents were capable of self-awareness to modify strategies based on their prior experiences. In addition to this, agents were capable of learning from other successful strategies, and were also capable of training others to increase knowledge of pest management. Despite the short-term costs incurred by implementing cooperative behaviour, model dynamics predicted decreased overall infestation levels over time.

Another example of this technique was utilized in a CHES model by Walsh et al. [117]. Using an ABM approach, a detailed model was developed to examine the relationship between an agricultural village population and their environment. Social modules were developed describing individual changes in social networks, population and assets, and land dynamics regardless of current climatic conditions.

## 4 Discussion

Our review shows that the number CHES models published each year has increased over the past decade. Moreover, there is a growing diversity in modelling techniques, including in the ways that human behaviour is represented. We found diverse methodologies in the construction of CHES models, including DEs (44/92), agent-based models (28/92) and stochastic models (9/92), with the remainder of the literature labelled as ‘other’ (11/92). With respect to modelling the human component, most papers used either optimization techniques (28/92) or modelled social influences with (34/92) or without (22/92) a game-theoretic foundation. Model complexity varied considerably. For instance, some papers using computational programs such as STELLA, or FSM [78, 7] calibrated their models using numerous parameter values generated from various data sources. In some cases, these models have also been able to successfully replicate historical trends [103].

We observed striking differences in the degree to which different environmental and ecological systems were represented in the literature: some types of natural systems were explored much more than other types. As observed in Table 2, CHES models have been implemented most commonly in agricultural settings, or in generic system models. Water allocation and tourism were the focus of the fewest papers. Furthermore, the impact of anthropogenic stress on the dynamics of terrestrial wildlife appeared to be relatively unexplored. Given that both marine and grassland systems are subject to invasive species and direct exploitation by humans, CHES modelling techniques could be applied to terrestrial wildlife dynamics more widely than they presently are. As an example, modelling techniques used to study overfishing of marine resources can be utilized as a template to build models for studying the dynamics of human-wildlife interactions with respect to hunting and poaching, especially with regard to restoration of critically endangered species. We speculate that the variation in how often different types of systems were studied over the past decade has more to do with ‘leader effects’ in the process of selecting research study systems, rather than inherent suitability or desirability of systems for CHES modelling.

Another topic that has not been explored in great detail is the effect of tourism on elements other than marine ecosystems. Touristic aggression on ecosystem dynamics have not been investigated in great detail in forest or grassland systems, despite the pervasiveness of tourism in those systems. As such, there is a great opportunity to study ecosystem interference and stress caused by tourism in a CHES framework.

As mentioned previously, the number of published CHES models increased significantly over the past decade. One possible reason for this increase could be diminishing computational barriers to developing complicated models, both with respect to computational power as well as making modelling accessible through specialized software like STELLA. As computational power has grown, it has become easier both to simulate complex models and parameterize them in increasingly sophisticated ways. Increasing complexity, on average reflects a general trend in model development in many fields, where models begin simple and over time become enhanced and increasingly complex [77]. Progression is due to the fact that simple models require less time to analyze, and provide insight on the opportunities to explore and enhance their explanatory power. In the context of CHES, these can take the form of newly recognized opportunities to expand existing ecological or environmental models to accommodate the increasing evidence of human impact and human inter-relatedness with natural systems. This does not imply that simple models are necessarily less accurate, but rather that they can provide the foundation for more complex future iterations based on the critical assumptions made during their initial development.

Misinformation is an important determinant of decision-making [107]. Most of the models we reviewed did not address misinformation explicitly. Many papers from outside our included literature are concerned with the spread of fake news and how to stop it, using a range of modelling techniques [109, 22, 43]. One paper employed approaches ranging from stochastic models to difference equations [22] to study the spread of fake news. Another example used game theory to investigate the Nash equilibrium strategy of a hypothetical ‘digital’ citizen who combats fake news regarding COVID-19 [43], while others have used a network model used to simulate the spread of hoaxes and their debunking [109]. However, some of the models included in our review studied imperfect information (which differs from misinformation in that the latter is deliberately falsified). In imitation dynamics models, for instance, it is often assumed that individuals respond to some proxy of ecosystem health (such as perceived forest cover or perceived climate events) rather than acting in full knowledge of current or future impacts on the natural system [12, 25].

Human interactions have evolved over the past few centuries from predominantly in-person interactions to communication at a distance through telephones and, more recently, online social media. With the introduction of online forums, groups and various outlets of social media, individuals can be exposed to different individuals with different beliefs, strategies and cultural norms [124, 4, 29].

Overall, these changes in the scale over which social forces operate will change individual relationships to natural systems. Moreover, we believe that the data provided by these sources will provide great insight into how individuals make decisions concerning natural systems, and how they are influenced by social forces.

Emerging data from new study systems suggests both new topics for behavioural modelling as well as new modelling techniques. For instance, the advent of online social media has led to the formation of social groups that are harmful to their own members, such as pro-anorexia groups [37]. The combination of changes to social conformity (e.g. peer pressure) and new digital means for socializing [33, 27, 123], mean that individuals can meet and interact with various others who share similar or different ideologies. Based on the guidelines of these groups, one must conform to a strict set of rules in order to be accepted. Thus, an individual using alternative strategies can be sufficiently influenced to adopt the beliefs set by the online group. In summary, the definition of one’s community interactions is not limited to strictly interpersonal interactions, and as such mathematical models can be extended to accommodate online influence in social networks. Such situations can be approached using social network simulation models.

In conjunction with the growth of online communities, there also appear to be opportunities for using simulated environments to observe human interactions in response to various stresses. An example of this occurred in an accidental epidemic caused by Blizzard Entertainment in their MMORPG (massively multiplayer online role playing game) World of Warcraft, where their player-base reacted to a rampant in-game epidemic [67]. This represents a natural experiment that is not dissimilar to controlled experiments used in economic game theory, for instance [18, 120]. Using simulated systems such as these, alternative data for measuring complex psychological behaviour can be obtained. For study populations where insufficient empirical data are available, trends from simulated environments can be extrapolated into CHES models to achieve a data-driven representation of human-environment interactions and potential intervention scenarios that can be explored.

Coupled human-environment systems models can hasten the identification of the forces that most strongly impact natural systems in the Anthropocene Era, and improve the representation of natural systems mathematical models. The literature we collected represents diverse and numerous variations of CHES modelling with similar long-term goals of not only explaining empirical observations, but also achieving sustainable trajectories for ecosystem dynamics. Continuing improvements in computational power and accessibility of model development software, together with growing data on both natural and human systems in the era of widespread digital data, suggest CHES modelling will continue to grow as a field and become more useful for applications to finding pathways toward environmental sustainability.ctories of environmental sustainability.

